# Transcribed microsatellite allele lengths are often correlated with gene expression levels in natural sunflower populations

**DOI:** 10.1101/339903

**Authors:** Chathurani Ranathunge, Gregory L. Wheeler, Melody E. Chimahusky, Andy D. Perkins, Sreepriya Pramod, Mark. E. Welch

## Abstract

Microsatellites are common in most species. While an adaptive role for these highly mutable regions has been considered, little is known concerning their contribution towards phenotypic variation. We used populations of the common sunflower (*Helianthus annuus*) at two latitudes to quantify the effect of microsatellite allele length on phenotype at the level of gene expression. We conducted a common garden experiment with seed collected from sunflower populations in Kansas and Oklahoma followed by an RNA-Seq experiment on 95 individuals. The effect of microsatellite allele length on gene expression was assessed across 3325 microsatellites that could be consistently scored. Our study revealed 479 microsatellites at which allele length significantly correlates with gene expression (eSTRs). When irregular allele sizes not conforming to the motif length were removed, the number of eSTRs rose to 2379. The percentage of variation in gene expression explained by eSTRs ranged from 1–86% when controlling for population and allele-by-population interaction effects at the 479 eSTRs. Of these, 70.4% are in untranslated regions (UTRs). A Gene Ontology (GO) analysis revealed that eSTRs are significantly enriched for GO terms associated with cis- and trans-regulatory processes. These findings suggest that a substantial number of transcribed microsatellites can influence gene expression.

## INTRODUCTION

The molecular basis of adaptation is of fundamental concern to evolutionary geneticists. Adaptation can occur via selection acting on standing genetic variation, or by that acting on novel mutations. While standing genetic variation might allow populations to respond rapidly to novel selective pressures in the short-term, novel beneficial mutations may arise at a finite rate. The relative contributions of these two processes have been argued (Orr 2010). What has been particularly puzzling to many theorists is the abundance of genetic variation in natural populations despite the expectation that its loss due to selection and drift should eliminate it. Ultimately, limited availability of heritable variation should serve as a form of evolutionary constraint. In spite of these factors that should limit the availability of additive genetic variation, there is much evidence to suggest that numerous traits can continue to respond to selection even after many generations of intense directional selection (Dudley and Lambert 1992; Mackay 1995; Yoo et al. 1980). These empirical results are consistent with the existence of mechanisms that continuously generate additive genetic variation at rates commensurate with its loss through selection.

These observations led Barton (1990) to suggest the existence of an abundant source of non-deleterious mutations that could affect quantitative traits. Kashi et al (1997) further emphasized that to be considered advantageous mutators with significant contributions toward rapid adaption, these mutations should be widely dispersed across genomes, associated with functional regions as regulatory elements or function as components of coding regions, and undergo high mutation rates leading to quantitative effects on phenotypes. These suggested favorable features are consistent with those observed in highly mutable microsatellites.

Microsatellites, or simple tandem repeats (STRs), are genomic regions consisting of short motifs that are 1- 6 bp in length, tandemly repeated up to a few dozen times (Vogt 1990). Microsatellites show high indel mutation rates (10^−2^ −10^−6^ / generation) (Ellegren 2004) resulting from mechanisms that may include replication slippage (Tautz and Renz 1984). Their mutation rates are estimated to be several orders of magnitude greater than that observed in non-repetitive DNA (Jarne and Lagoda 1996; Li 1997). Despite their apparent fit for the role of advantageous mutators, microsatellites have long been perceived as neutrally evolving, non-functional regions, and have been used as the “molecular marker of choice” in population genetics and forensics (Jarne and Lagoda 1996). This long-standing perception was questioned by the abundance of highly mutable microsatellites in structural regions placing them in positions that could influence gene function and gene products (Li et al. 2002; Li et al. 2004). Further, non-random distribution of microsatellites in genomes has been reported in several organisms, including fruit fly (*Drosophila melanogaster)* (Bachtrog et al. 1999), thale cress (*Arabidopsis thaliana)*, rice *(Oryza sativa*) (Lawson and Zhang 2006; Morgante et al. 2002), and common sunflower (*Helianthus annuus* L.) (Pramod et al. 2014). Microsatellite variation in functional regions has been consequently linked to phenotypic variation. Notable examples include microsatellites linked to human neurodegenerative diseases such as fragile X syndrome (Verkerk et al. 1991) and Huntington’s disease (Andrew et al. 1993).

The claim that microsatellites can generate adaptive genetic variation is now supported by a growing list of studies. Research has linked microsatellites to variation in skeletal morphology in domesticated dogs (Fondon and Garner 2004), social behavior changes in voles (Hammock and Young 2005) and some primates (Hopkins et al. 2012), pathogenesis in bacteria (Moxon et al. 1994; Moxon et al. 2006), plasticity in adherence to substrates in *Saccharomyces cerevisae* (Verstrepen et al. 2005), and thermal sensitivity in *Drosaphila melonogaster* (Costa et al. 1991) among others. However, little is known about the mechanisms by which microsatellites can influence phenotypes. An intriguing model proposed to explain the underlying mechanisms by which microsatellites may influence phenotypes is the “tuning knob” model. This model likens microsatellites to tuning knobs where stepwise changes in microsatellite allele lengths can have stepwise effects on phenotypes, allowing selection to adjust or fine tune a population’s phenotypes corresponding to environmental stresses (Kashi and King 2006; King et al. 1997; Trifonov 2004). The model explains that these effects can be mediated either by modulating gene expression or by facilitating structural changes in proteins.

Microsatellites located upstream of genes as well as those in 5’ untranslated regions (UTRs) and introns might affect gene expression while those in 3’ UTRs are more likely to influence transcript stability (Li et al. 2004). Microsatellites located in coding regions are likely to generate structural changes in proteins, but may also play a regulatory role as well (Gemayel et al. 2010; Li et al. 2004). The role of microsatellites in modulating gene expression has been supported empirically by introducing microsatellites of different lengths to promoter regions of genes in *Saccharomyces cerevisiae* (Lee and Maheshri 2012; Vinces et al. 2009) and in Tausch’s goatgrass (*Aegilops tauschii*) (Ryan et al. 2010). Experimentally introduced microsatellite tracts in coding regions of genes have also been utilized to show their role in altering protein structures in *Arabidopsis thaliana* (Golubov et al. 2010). It is apparent that many studies have focused on the effect of microsatellites on specific genes, as opposed to investigating the global role that microsatellites may play across genomes potentially leading to adaptive evolution. Such large-scale genome level studies focusing on potential phenotypic effects of multiple microsatellite loci across populations have been few and sporadic (Fahima et al. 2002; Gymrek et al. 2015; Nevo et al. 2005). Here, we attempt to test the relevance of the tuning knob model at the genome level using transcribed sequences. Microsatellites located within transcribed regions are particularly accessible in this regard given that RNA-Seq data can be used to both genotype transcribed microsatellites, and allow for estimation of allele-specific expression at microsatellite-encoding transcripts.

In this study we use natural populations of the common sunflower (*Helianthus annuus* L.) transecting a latitudinal cline from Kansas to Oklahoma. We chose sunflower as a model system given their adaptability to diverse environmental conditions across their broad geographical range in North America (Heiser et al. 1969). Particularly, populations across latitudes demonstrate heritable clinal variation in a number of traits including flowering time (Blackman et al. 2011) and seed oil content (Linder 2000). These distinct patterns of variation in adaptive traits across latitudinal populations provide an opportunity to test the hypothesis that microsatellites generate some of the adaptive genetic variation found within and among natural populations.

From a probabilistic viewpoint, it is reasonable to expect that transcribed microsatellites, due to their proximity to functional regions, are likely to influence gene expression as *cis-*regulatory elements. We utilized an RNA-Seq approach to assess the degree to which allele lengths of transcribed microsatellites influence gene expression across multiple transcripts to identify potential tuning knobs. We predicted that allele lengths of transcribed microsatellites that function as tuning knobs are correlated with gene expression levels.

## MATERIALS AND METHODS

### Sample collection and common garden experiment

As previously described, seeds were collected from six natural populations of common sunflowers from two latitudinal locations in Kansas and Oklahoma (three populations from each location) (Supplemental Table S1) (Ranathunge *et al*. 2018). Scarified seed were germinated on moist filter paper in petri dishes. Seedlings were transferred into 2.54 cm “cone-tainers” (Stuwe & Sons, Inc., Tangent, OR, USA) arranged in a randomized design and were kept in a greenhouse under controlled conditions. At the age of four weeks, young leaf tissue samples from 96 individuals representing the 6 populations (16 individuals from each) were collected for RNA isolation. Multiple populations from each latitude were used so that the relative effect of the microsatellite allele length on gene expression could be assessed while controlling for other components of phenotypic variation such as variation in the environment and the local genetic background.

### RNA-Seq and de novo transcriptome assembly

RNA was isolated from 20 mg of fresh leaf tissue with Maxwell 16 LEV *simply*RNA Tissue kits (Promega, WI, USA). Isolated RNA samples were sent to HudsonAlpha Institute for Biotechnology (http://hudsonalpha.org) for high throughput sequencing. The procedure for cDNA library preparation and RNA sequencing was described in detail in Ranathunge *et al* (2018). RNA sequencing (2 × 100 paired end) was carried out with Illumina HiSeq 2500 platform. High quality reads were obtained for 95 out of 96 individuals. The average total number of reads per individual was 42.8 million. The total number of reads per individual ranged from 27.6 – 88.1 million (Supplemental Table S2). RNA-Seq produced 100 bp paired end reads. The quality of the reads was assessed with Fast QC software (http://www.bioinformatics.babraham.ac.uk/projects/fastqc/) using default settings. A deviation from expected base frequencies was observed at the beginning of sequenced fragments, indicating possible adapter contamination; these positions were removed from the reads by trimming 14 bp from each end. Trimmed reads complying with the default quality standards of the FastQC software were used in the downstream processes. The sample that produced the highest total number of reads (88.1 million) which indicated the best coverage and also complied with base quality standards was used to construct a reference transcriptome. The reference transcriptome was built with the software modules (Inchworm, Chrysalis, and Butterfly) of the Trinity program (Grabherr et al. 2011). We used the paired end option in Trinity with a minimum contig size set to 200 bp and the minimum k-mer coverage set to 1. The reference transcriptome consisted of 58,431 contigs. Fastq files with raw reads obtained from the remaining 94 individuals were aligned to the reference transcriptome with Bowtie2 (Langmead and Salzberg 2012). Bowtie2 produced contigs were managed using SAMtools (Li et al. 2009). SAMtools produced BAM format files which were then sorted and indexed to quantify gene expression. Gene expression was quantified from reads aligned to the reference transcriptome. The reads for each contig were normalized by the library size of each individual, and gene expression was estimated as reads per 100 million mapped reads as explained in Ranathunge et al. 2018.

### Functional annotation

A standalone BLAST (Basic Local Alignment Search Tool) search (Altschul et al. 1997) was performed on the reference transcriptome against the *Helianthus annuus* protein sequence database to annotate the reference transcriptome. First, protein sequence data for sunflower unigenes were downloaded from the sunflower genome database at https://www.sunflowergenome.org/ and a database was created. Sequences from the reference transcriptome were used as the query in the BLASTX search. An E-value cutoff of 0.0001, gap open penalty score of 11, gap extension penalty score of one and minimum word size of three were used as alignment parameters. BLOSUM62 was used as the matrix of choice. A best hit overhang of 0.25 and maximum target sequence value of one were used to minimize the number of hits for each query sequence. The results of the BLASTX search were output in tabular format and the hits were further filtered based on bit score and e-value to retain the single hit with the highest bit score and lowest E-value for each query sequence.

### Microsatellite genotyping

We searched the reference transcriptome for microsatellites with Tandem Repeat Finder (Benson 1999). The match, mismatch, indel and alignment scores were set at 2, 7, 7, and 50 respectively. The parsed output file from Tandem Repeat Finder and BAM format files from the 95 individuals were used as input for RepeatSeq (Highnam et al. 2013) for genotyping microsatellites. RepeatSeq uses a Bayesian approach for making variant calls on reads generated with short-read sequencing technology. RepeatSeq’s default settings were used to genotype transcribed microsatellites across all individuals. The output from RepeatSeq was initially used to assess the motif size and type frequencies within the list of genotyped microsatellites. Motif types were standardized with custom PERL scripts as per the motif matrix mentioned in Kofler et al (2007). Output files from RepeatSeq were also used to extract position of the microsatellite in the transcript.

Reads that failed to generate genotypes were removed from the downstream processes. Raw expression estimates based on read counts were obtained for each allele for each individual with data corresponding to the major component, allele genotype, reference genotype, and motif. We noticed that RepeatSeq tends to erroneously call certain genotypes as heterozygous by identifying underrepresented error reads as alleles. To solve this issue, we identified a genotype as heterozygous only when the second most represented allele was called with at least 25% of the frequency of the dominant allele. In the majority of cases with more than two alleles present, many were at very low frequency and were removed by this method.

### Effect of microsatellite allele length on gene expression

We examined the effect of microsatellite allele length on gene expression (log-transformed) by employing an analysis of covariance (ANCOVA) while controlling for population and allele-by-population interaction effects. Linear, quadratic, and cubic regression were all attempted because previous *in-vivo* experiments have identified non-linear relationships between microsatellite allele length and gene expression (Vinces et al. 2009). A p-value of 0.05, and q-value cutoff of 0.05 that corrects for multiple comparisons (Storey 2002), were used to identify microsatellites where allele length correlates with gene expression (hereinafter referred to as eSTRs - a term borrowed from Gymrek et al. 2015 which refers to significant expression short tandem repeats). All statistical analyses were performed in R version 3.3.3 (R Core Team 2017). To identify eSTRs, we first employed a conservative approach by conducting ANCOVAs with unfiltered microsatellite allele lengths and allele specific gene expression estimates. This first analysis included irregular allele sizes inconsistent with the length of the repeat motif. We then removed these alleles from the analyses and reran the ANCOVA to estimate the microsatellite allele length effect on allele-specific gene expression.

Further, information from the RepeatSeq output was used to calculate motif size and type frequencies within eSTRs to assess whether microsatellites of specific motif size and type classes are more likely to function as tuning knobs. We used the BLASTX output for the reference transcriptome to extract best hits for the eSTR-containing transcripts. Information on the reading frame, “query start” and “query end” positions from the BLASTX output along with eSTR position data from the RepeatSeq output were used to identify whether an eSTR was located within the 5’UTR (Untranslated region), coding region, or the 3’UTR. To test whether mechanisms underlying expansion and contraction of eSTRs are likely to be different based on the location of eSTRs and/or their respective motif sizes, we conducted Kruskal-Wallis (KW) tests on eSTR tract lengths. The KW tests statistically compared eSTR tract lengths among the three regions, 5’UTR, coding region and 3’UTR, and among eSTRs of different motif sizes.

### Validation of gene expression estimates by Real Time PCR (qPCR)

We conducted qPCR analysis on seven eSTRs selected based on the magnitude of allele length effect on gene expression to validate RNA-Seq based gene expression estimates. The isolated RNA samples from 48 of the individuals used in the RNA-Seq experiment were converted to cDNA with High Capacity cDNA Reverse Transcription kit with RNase inhibitor (Applied Biosystems, Foster City, California, USA). Sequence data were obtained for seven eSTR-containing transcripts from the *de novo* transcriptome. TaqMan gene expression assays were designed for the seven selected loci using Primer Express version 3.0 (Applied Biosystems, Foster City, California, USA) (Supplemental Table S3). TaqMan assays for two constitutively expressed genes in sunflower, actin and ubiquitin, previously designed by Pramod et al. (2012) were used as controls. Protocols for generating standard curves and validating RNA-Seq derived gene expression estimates are given in Supplemental Note 1 and Supplemental Table S4.

### Gene Ontology (GO) enrichment analysis

Protein sequences corresponding to the best hits for the reference transcriptome obtained from the BLASTX search were retrieved. The sequences were used to conduct a Gene Ontology (GO) analysis with Blast2GO (Conesa et al. 2005) to identify GO terms associated with all genes in the reference transcriptome. A BLAST search was conducted against the *Arabidopsis thaliana* protein sequence database with the blastp option. An E-value cutoff of 0.001, a minimum number of BLAST hits at 20, a HSP length cutoff of 33, and a word size of 3 were used as settings in the blastp search. The BLAST hits were then mapped and annotated with default settings to identify GO terms under three categories, namely, biological processes, molecular function, and cellular component. To identify specific GO terms enriched within eSTR-containing transcripts, Fisher’s exact test was performed with GO terms associated with all *H. annuus* genes in the reference transcriptome as the background list and *H.annuus* gene IDs corresponding to eSTR-containing transcripts as the test set. A False Discovery Rate (FDR) of 0.05 was used as the significance threshold to identify overrepresented GO terms within the eSTR-containing transcripts. General GO terms (parent terms) were removed to retrieve the most specific terms using “reduce to most specific” option in Blast2GO. The REVIGO (Supek et al. 2011) tool was also used to reduce the enriched GO terms to the most specific terms for visualization based on the uniqueness and dispensability scores calculated by the program.

## RESULTS

### Microsatellite search

Tandem Repeat Finder identified 19,104 potential microsatellites in the reference transcriptome. We were able to genotype 3,325 of these microsatellites in 2,640 transcripts consistently using RepeatSeq. A total of 685 (25.9%) transcripts harbored more than one microsatellite. Hexanucleotides (38.89%) were identified as the most abundant microsatellite motif length, followed by trinucleotides (29.17%) (Supplemental Table S5; Supplemental Fig.S1). A total of 355 standardized motif types were identified. Of the 355 different motif types identified, the ACC repeat motif has the highest frequency (6.32%), followed by ATC repeats (5.78%) (Supplemental Table S6; Supplemental Fig.S2).

### Microsatellites with significant allele length effect on gene expression (eSTRs)

When individuals with irregular allele sizes were included in the ANCOVA, 816 (24.5%) of the microsatellites scored showed a significant allele length effect on gene expression (log transformed) (p < 0.05), controlling for population and allele-by-population interaction effects. After correcting for multiple comparisons (Storey 2002), a total number of 479 (14.4%) microsatellites in 449 unique transcripts were identified as eSTRs (Supplemental Table S7). The percent of variation in gene expression explained by allele lengths ranged from 1% - 86% in the 479 eSTRs. ANCOVA results for 237 microsatellites suggest a positive relationship between allele length and gene expression levels while 242 microsatellites exhibited a negative correlation between allele length and gene expression (Figure 1). When the irregular allele sizes that did not conform to the motif length were removed from the analysis, the number of eSTRs rose to 2379 (71.5% of the genotyped microsatellites) after correcting for multiple comparisons (Storey 2002). The effect of allele length on gene expression ranged from 0.077 to 79.9% in the 2379 microsatellites (Supplemental Table S8). While we acknowledge that the first approach could have been conservative in identifying eSTRs, removal of all irregular allele sizes from the analysis could have resulted in excessive manipulation of data as demonstrated by the inflated estimates of correlation between allele length and gene expression. Taking these concerns into consideration, we used the 479 eSTRs identified with the first approach for downstream analyses.

**Figure 1.**
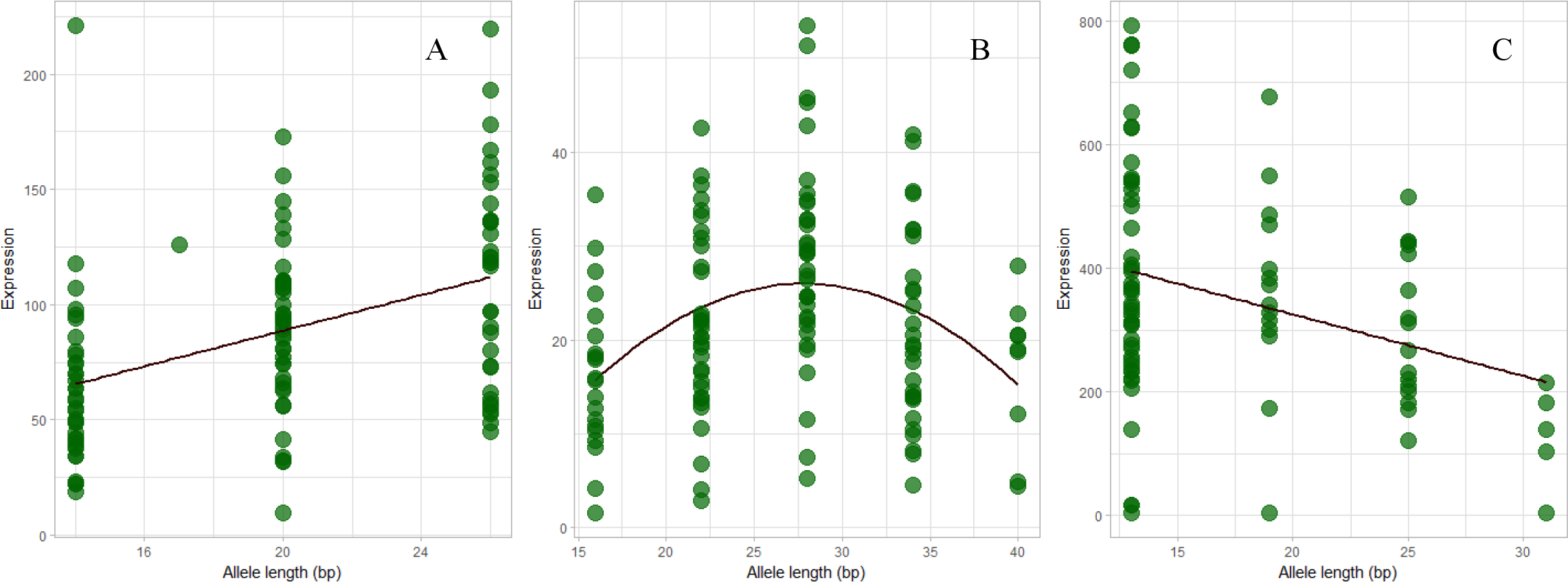
The effect of microsatellite allele length on gene expression. An Anlaysis of Covariance (ANCOVA) revealed different patterns of correlation between microsatellite allele length and allele specific gene expression (eSTR). (A), (B) and (C) show positive, quadratic and negative correlation patterns observed between allele length (bp) and allele specific gene expression (read counts per 100 million reads) in microsatellite loci located in contigs, comp26672 (CCTTCT), comp49389 (GAACCA) and comp25013 (GACGGT), respectively.

We characterized the motif lengths associated with eSTR-containing transcripts in addition to the location of the microsatellite relative to start and stop codons. Hexanucleotides (40.1%) are the most abundant motif length within the 479 eSTRs followed by trinucleotides (31.5%) (Figure 2; Supplemental Table S9). A total of 169 different motifs (standardized) were identified within 449 transcripts (0.38 motif/transcript) containing the 479 eSTRs. ACC (21.9%) was the most abundant motif within eSTRs followed by AAG (17.2%) (Figure 2; Supplemental Table S10). Prior work detected significant enrichment of A and AG motif-containing microsatellites within genes that are differentially expressed between the two latitudes in Kansas and Oklahoma (Ranathunge *et al*. 2018). This suggested that A and AG repeat motifs are likely to be involved in gene expression divergence among these sunflower populations. To test whether specific motif-containing microsatellites are more likely to function as tuning knobs for gene expression, we estimated enrichment of different microsatellite motif types within eSTRs compared to the remaining 2846 microsatellites genotyped (non-eSTRs). However, we did not detect any significant enrichment of specific motif types within eSTRs compared to non-eSTRs (Chi-squared test, p > 0.05).

**Figure 2.**
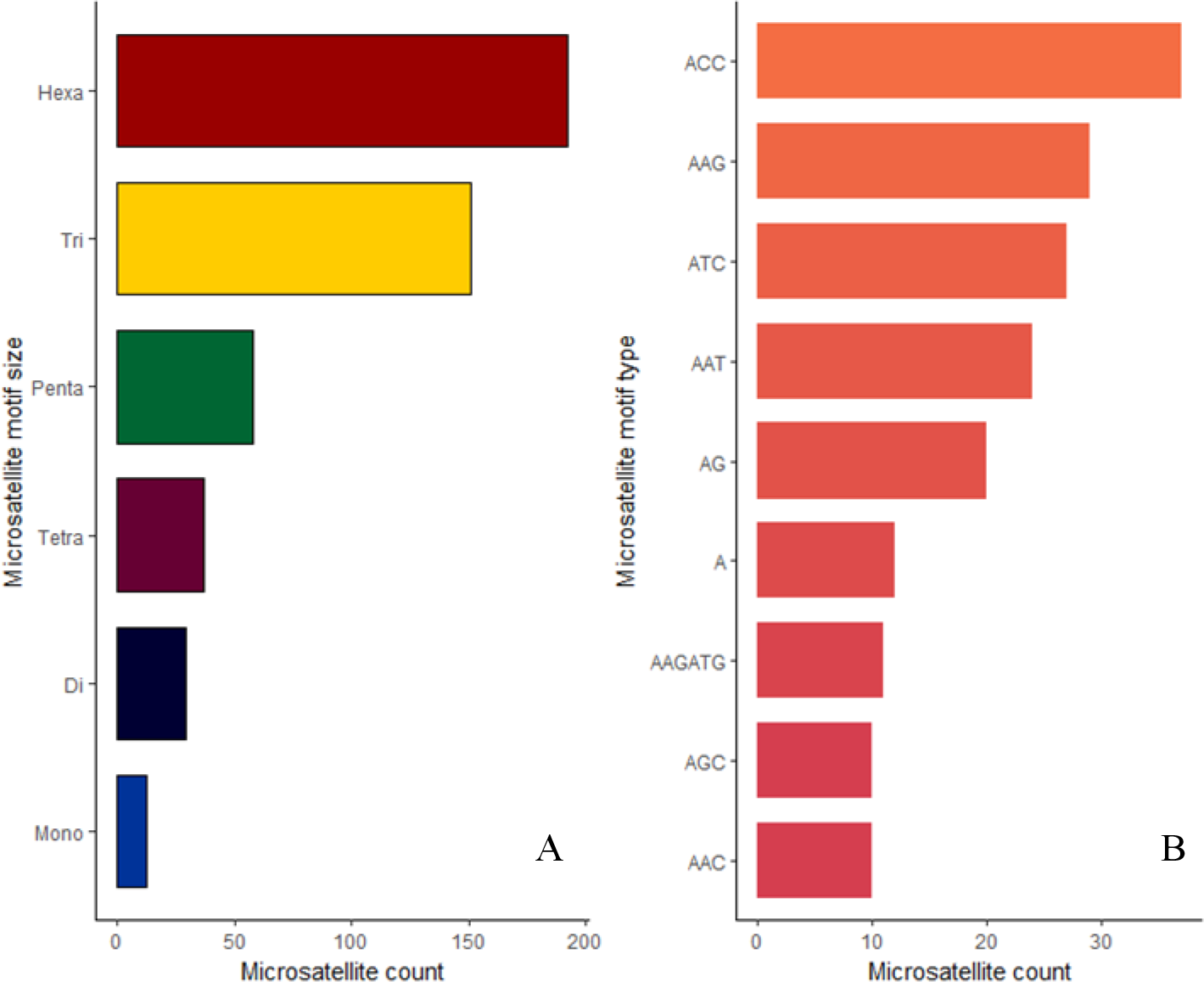
The distribution of microsatellite motif sizes and types in eSTRs. (A) Number of mono-, di-, tri-, tetra-, penta- and hexanucleotides identified within eSTRs. (B) The top nine microsatellite motif types and their counts within eSTRs.

eSTRs were most abundant within 5’UTRs (42.1%) followed by coding regions (29.6%) and 3’UTRs (28.3%) (Figure 3). Of the 165 eSTRs located within the 5’UTRs, the most abundant motif size present were trinucleotides (34.55%) followed by hexanucleotides (30.30%) (Figure 3). Within the 116 eSTRs in the coding regions, hexanucleotides were the most abundant motif size (56.03%) followed by trinucleotides (37.93%) while dinucleotides were absent in the region. Mononucleotides (0.86%), tetranucleotides (1.92%), and pentanucleotides (0.96%) were also scarce within coding regions (Figure 3). Hexanucleotides were also the most abundant within the 111 eSTRs located within the 3’UTRs (38.74%) followed by trinucleotides (26.13%) (Figure 3). ACC was the most abundant motif in all three regions; 5’UTR (8.48%), coding region (8.62%) and 3’UTR (8.11%) (Supplemental Table S11). Prior work revealed that a transcript containing a microsatellite within the 3’UTR is more likely to be differentially expressed between the latitudinal populations in Kansas and Oklahoma (Ranathunge et al. 2018). In line with this prediction, we tested whether a transcript containing a microsatellite within 5’UTR, coding, or 3’UTR is more likely to function as tuning knobs for gene expression. We assessed the enrichment of eSTRs within 5’UTRs, coding regions, and 3’UTRs in comparison to frequency of non-eSTRs within the three regions. We did not detect any significant enrichment of eSTRs within any of the three regions (Chi-squared test, p > 0.05). Further, there was no significant difference in mean microsatellite tract lengths among eSTRs located in 5’ UTRs, coding regions and 3’ UTRs (KW test, p = 0.45) (Figure 4). No significant differences were detected in eSTR tract length among different motif sizes (KW test, p = 0.07) (Figure 4).

**Figure 3.**
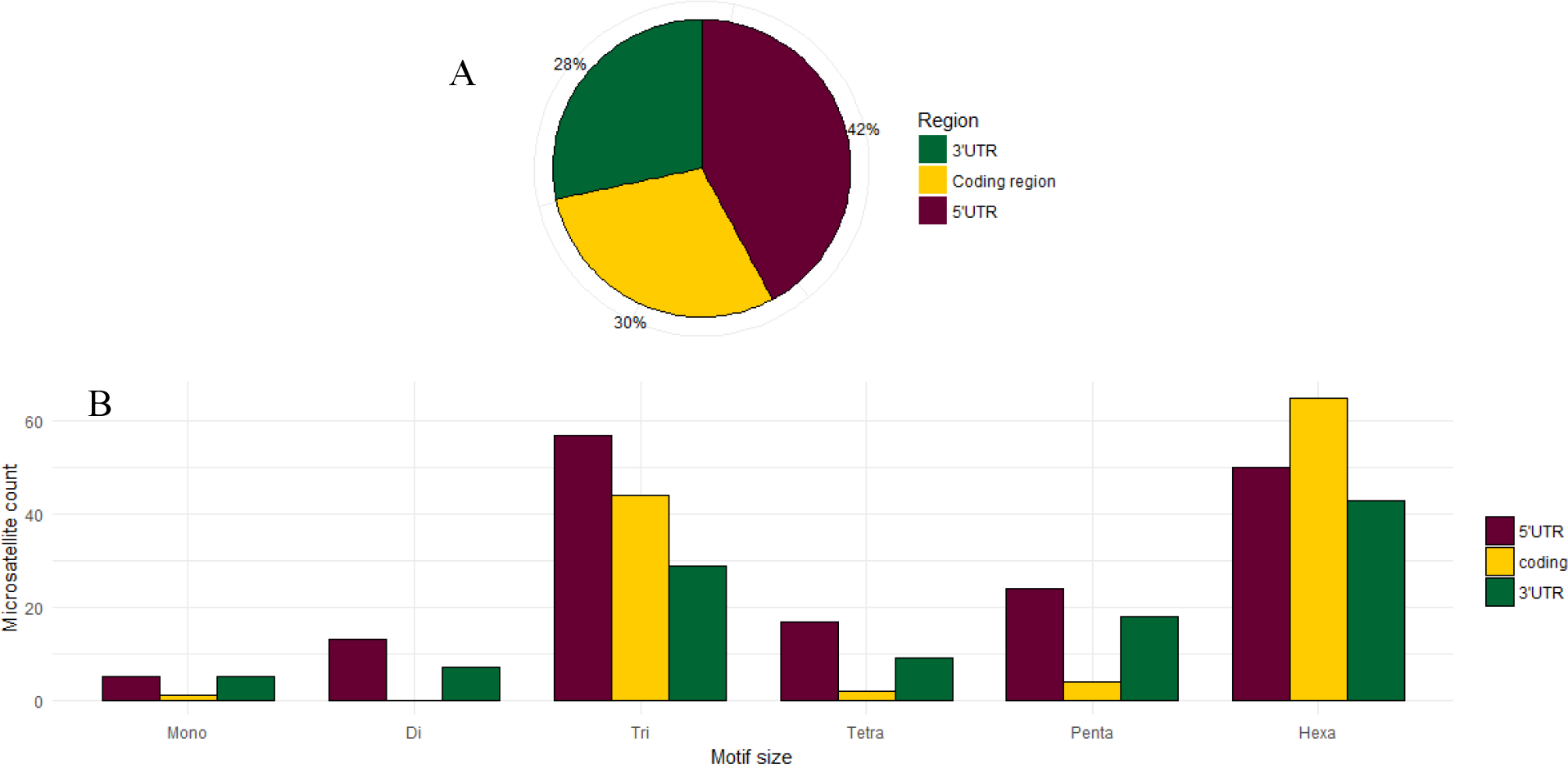
The distribution of eSTRs located within the three regions; 5’UTR, coding and 3’UTR. (A) Pie chart represents the percentage of eSTRs located within the three regions; 5’UTR, coding and 3’UTR. (B) The counts of different eSTR motif sizes identified within each region.

**Figure 4.**
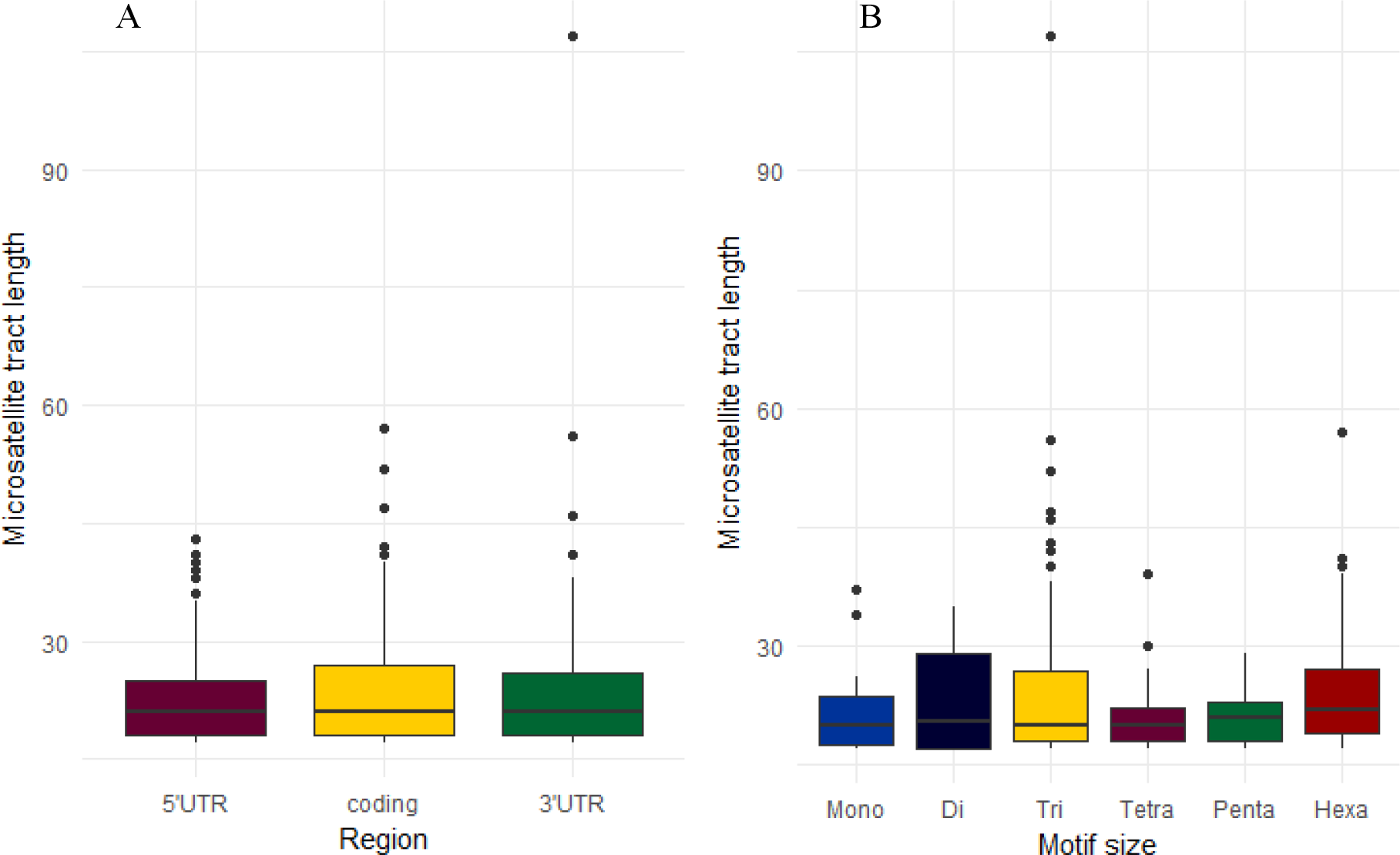
The variation in eSTR tract lengths. (A) The variation in eSTR tract length by region. (B) The variation in eSTR tract length by motif size.

### Validation of gene expression estimates by Real Time PCR (qPCR)

All seven qPCR assays for eSTR-containing transcripts showed strong log-linear relationships between C_T_ values and cDNA concentrations. Coefficient of determination estimates (R^2^) ranged from 0.97 – 0.98 and efficiencies (b) ranged from 0.71 to 0.9 (Supplemental Table S4). Regression analyses conducted with log C: R ratios for pairs of high copy number and low copy number loci revealed strong correlation between loci for relative concentrations estimated with qPCR and RNA-Seq data with R^2^ values ranging from 0.77 to 0.96 (Supplemental Table S12; Supplemental Fig.S3).

### Functional annotation and Gene Ontology (GO) enrichment analysis

The BLASTX search against the *H. annuus* protein database identified unique hits for 17,985 contigs in the reference transcriptome (Supplemental Table S13). Of the 449 eSTR-containing transcripts, 371 had blast hits that passed the significance threshold (Supplemental Table S14). In comparison to GO terms associated with the annotated 17,985 genes in the reference transcriptome, eSTR-containing transcripts were significantly enriched for 57 GO terms (Supplemental Table S15). The simplified enriched GO term list included eight specific GO terms (Table 1; Figure 5). They were classified under biological process (4), molecular function (2) and cellular component (2). The most enriched GO term within the eSTR-containing transcripts was associated with regulation of transcription (GO:0006355) in the biological process category (Table 1; Figure 5). Within the molecular function category, the GO term represented by most sequences in our list was transcription factor activity (GO:0003700) while the GO term, spliceosomal complex (GO:005681) was the most overrepresented within the cellular component category.

**Table 1.** Gene Ontology (GO) terms enriched within eSTR-containing transcripts in 373 *Helianthus annuus* (reduced to most specific terms)

| GO ID | GO Name | GO Category | FDR | p - value | % in test set | % in background set |
| --- | --- | --- | --- | --- | --- | --- |
| GO:0006355 | regulation of transcription, DNA-templated | BP | 6.93E-05 | 1.06E-07 | 17.73% | 8.66% |
| GO:0003700 | transcription factor activity, sequence-specific DNA binding | MF | 0.01169 | 5.86E-05 | 10.47% | 5.12% |
| GO:0003677 | DNA binding | MF | 0.024173 | 0.000129 | 15.12% | 8.84% |
| GO:0005681 | spliceosomal complex | CC | 0.025453 | 0.000145 | 2.33% | 0.41% |
| GO:0000398 | mRNA splicing, via spliceosome | BP | 0.027348 | 0.000158 | 2.62% | 0.53% |
| GO:0051347 | positive regulation of transferase activity | BP | 0.034644 | 0.000204 | 1.45% | 0.14% |
| GO:2000113 | negative regulation of cellular macromolecule biosynthetic process | BP | 0.035718 | 0.000215 | 6.69% | 2.85% |
| GO:0097526 | spliceosomal tri-snRNP complex | CC | 0.037228 | 0.000232 | 1.16% | 0.07% |

Note: BP = Biological process, MF = Molecular function, CC = Cellular component

**Figure 5.**
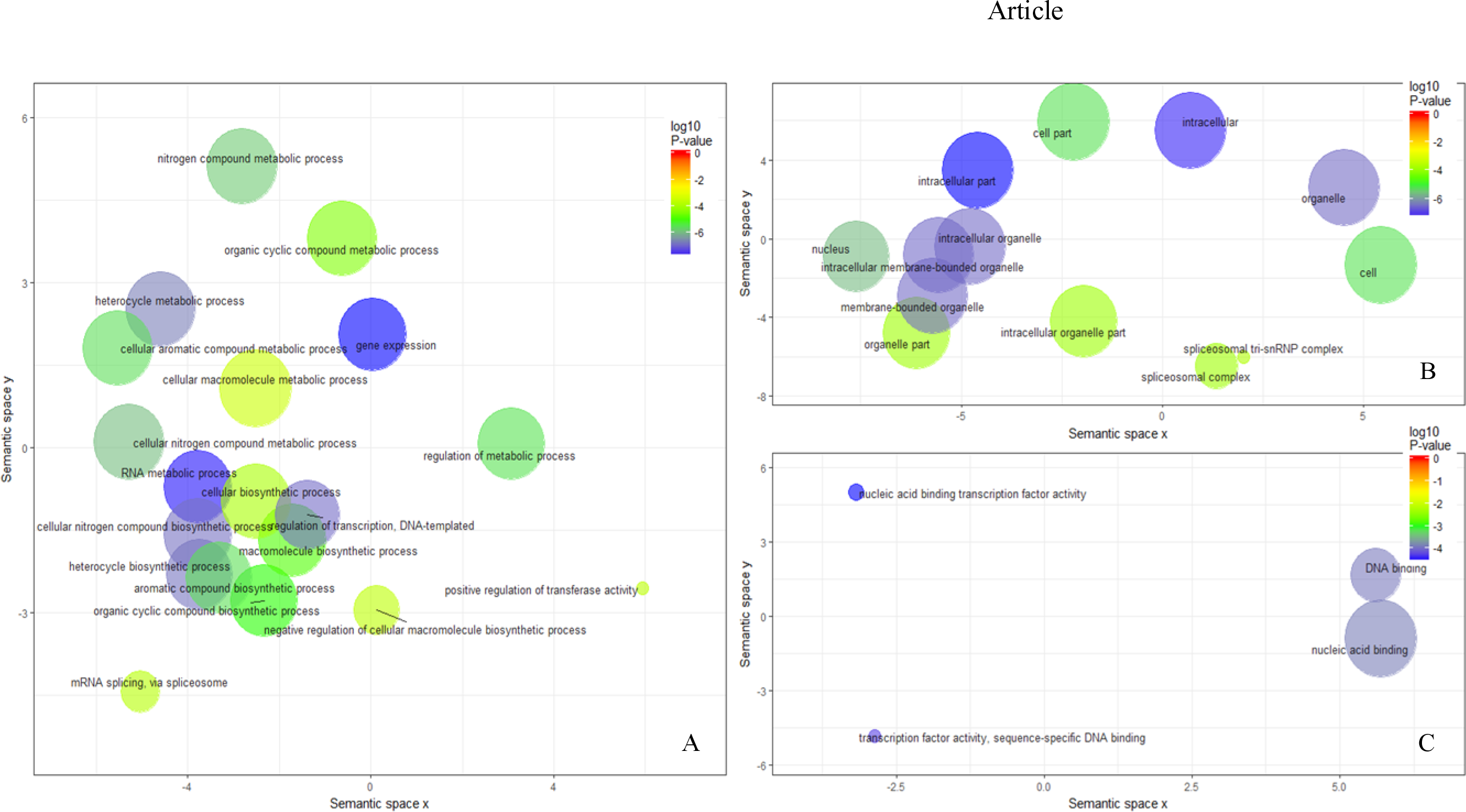
Gene Ontology (GO) terms enriched within eSTRs. GO enrichment analysis of eSTR-containing transcripts was conducted against a background of all expressed transcripts. The enriched terms were visualized using REVIGO. (A). Reduced GO terms related to biological processes. (B) Reduced GO terms associated with cellular components. (C) Reduced GO terms associated with molecular functions. The circle size represents the frequency of the GO term and the color represents the log_10_ P-value based on the Fisher’s exact test for enrichment.

## DISCUSSION

Here we tested specific predictions of the tuning knob model that proposes stepwise effects of microsatellite allele length on phenotypes. This study focused on transcribed microsatellites because both genotypic and phenotypic variation at the level of gene expression can be assessed using RNA-Seq data. We used *Helianthus annuus*, common sunflower, seed collected from two latitudes to infer the scope of this proposed mechanism for rapid evolutionary change in natural populations. The seeds were germinated, and plants were grown in a common garden experiment to minimize environmental variation. Further, we employed an experimental design that included replicates from each of the two latitudes which allowed us to estimate the relative effects of the microsatellite allele length on gene expression given the effects of other components such as local genetic background and environmental variation within latitudes on phenotypic variation. These populations also show latitudinal variation in a number of morphological traits including flowering time, and plant height at budding that indicates the heritable nature of phenotypic traits in these populations (Ranathunge *et al*. 2018). These observations provide an impetus to study the underlying mechanisms of adaptation across this latitudinal cline.

Using the RNA-Seq data from these populations, we were able to both estimate gene expression levels and consistently genotype microsatellites in 2,640 transcripts containing 3325 microsatellites. Motif type and length searches across the 3325 microsatellites showed frequency estimates consistent with similar surveys conducted on eukaryotic genomes (Qin et al. 2015; Tóth et al. 2000) including those conducted on the *Helianthus annuus* transcriptome with the use of an EST database (Pramod et al. 2014). These results demonstrate that the microsatellites used to test the predictions of the tuning knob hypothesis in this study substantially represent the composition of microsatellites in eukaryotic genomes (Supplemental Table S5; Supplemental Table S6).

Our findings from the ANCOVAs estimating the effect of microsatellite allele length on gene expression provide support for the tuning knob model at a large number of microsatellite encoding transcripts. Of the loci scored for both gene expression level and microsatellite genotypes (3325), 479 microsatellite loci (14.4%) (eSTRs) showed significant allele length effect on gene expression. It is important to note that this first analysis was conducted without filtering out irregular allele sizes that did not conform to the repeat motif. When irregular alleles were removed from the analysis, the number of eSTRs rose to 2379 (71.5%). The irregular allele lengths observed in some individuals could represent imperfect microsatellites resulting from substitutions and indels within the microsatellite tract or sequencing artifacts. Typically, imperfect microsatellites have been noted for lower mutation rates and higher stability in comparison to perfect microsatellites (Kunkel and Soni, 1988). Sequence interruptions have been previously reported in microsatellites linked to trinucleotide-repeat disorders (Chung *et al*. 1993; Eichler *et al*. 1994; Kunst *et al*. 1997). Based on these findings, it is reasonable to expect that mechanisms underlying functional microsatellites are likely to be affected by the level of stability attributed to these repeat tracts. Perhaps irregularities in microsatellites can hinder the cis-regulatory activity of these genetic elements.

Downstream analyses were limited to the 479 eSTRs identified with the more conservative of the two approaches. The 479 eSTRs showed on average, 1% – 86% allele length effect on gene expression when accounted for population and allele-by-population interaction effects. The majority of the eSTRs identified in ours study showed a linear relationship between microsatellite allele length and gene expression. Previous studies have also demonstrated positive and negative linear relationships between microsatellite length and gene expression (Contente et al. 2002; Gymrek et al. 2015; Shimajiri et al. 1999). We also tested whether our data fit a quadratic model as previous experimental studies have demonstrated similar patterns of correlation between experimentally constructed promoter microsatellite lengths and gene expression in yeast (Vinces et al. 2009). Based on the significance threshold we set when conducting the ANCOVA, 171 eSTRs showed support for a quadratic relationship between microsatellite allele length and gene expression. However, when we examined the data for those loci, most of these trends were identified as artifacts of rare allele lengths represented in the data set. These findings suggest that the relationship between gene expression and extant alleles in natural populations can typically be modeled as linear relationships.

The position of the microsatellite tract within genes may also provide insights regarding potential mechanisms by which microsatellites modulate gene expression. To better understand these different mechanisms, we estimated the frequencies of different motif types within the eSTRs, and also examined the likely locations for the eSTRs within *H. annuus* genes. A previous study conducted on the RNA-Seq data generated from the same populations indicated that microsatellites with motif types A and AG as well as microsatellites located within the 3’UTRs are likely to cause gene expression divergence among these latitudinal populations (Ranathunge et al. 2018). In the current study we did not detect any significant enrichment of specific microsatellite motif types within the eSTRs in comparison to non-eSTRs which suggests that the motif type may not affect the microsatellite’s ability to function a tuning knob. These contrasting patterns observed between the study on microsatellites involved in differential gene expression (Ranathunge *et al*. 2018) and the current study on eSTRs suggests the involvement of different groups of microsatellites in generating large and subtle differences in gene expression. Furthermore, except for one microsatellite locus in transcript comp29399, there were no microsatellites in common between those identified as eSTRs in the current study and the microsatellites identified within differentially expressed genes between these two latitudinal populations by Ranathunge et al. 2018.

When we examined the location of the eSTRs within the transcribed region, our results suggest that most eSTRs (42.1%) are located in 5’UTRs. Combined, microsatellites in 5’ and 3’ UTRs accounted for 70.4% of the eSTRs. However, when we estimated the enrichment of eSTRs in the three regions in comparison to the distribution of non-eSTRs genotyped, we did not detect any significant difference. This suggests that the likelihood of microsatellites functioning as tuning knobs may not depend on their location within the transcribed region. Several experimental studies have demonstrated that some microsatellites in 5’UTRs are vital for the expression of the gene (Kumar and Bhatia 2016; Streelman and Kocher 2002; Toutenhoofd et al. 1998). Some of the proposed mechanisms by which 5’UTR microsatellites may function as cis-regulatory elements involve serving as transcription factor binding sites (Kumar and Bhatia 2016). Microsatellites in coding regions may also play a role in gene expression regulation. In our study, 29.6% of the eSTRs were located in coding regions. Variation in coding region microsatellites is linked to changes in the structure of proteins including transcription factors as documented in several studies (Fondon and Garner 2004; Gemayel et al. 2015; Lee et al. 2003). However, the mechanisms by which they may function as cis-regulatory elements directly influencing gene expression levels are not entirely clear. Some triplet repeats including those in coding regions are proposed to affect nucleosome binding thereby affecting transcription rates (Sandman and Reeve 1999; Wang 2007). Coding region eSTRs identified in this study are well represented with triplet repeats that could potentially function in the same manner. The substantial percentage of coding region eSTRs identified in this study may also indicate previously unreported mechanisms by which they may function as cis-regulatory elements. Microsatellites in 3’UTRs are assumed to influence transcript stability via AU rich repeats that influence mRNA decay (Mignone et al. 2002). Mononucleotides in 3’UTRs have been proposed to play a role in regulating gene expression in number of cancer-related genes (Paun et al. 2009; Ziqiang et al. 2009). Rattenbacher et al.(2010) reported that GU rich elements in 3’UTRs can cause mRNA decay. Collectively, these proposed mechanisms may explain how specific eSTRs in 3’UTRs identified in this study may influence gene expression levels.

Tract length comparisons among eSTRs from the three regions did not indicate any significant difference suggesting that eSTRs may expand or contract under similar selective pressures irrespective of the region they are located. This finding runs contrary to previous studies that report significant tract length differences in general among microsatellites located in different regions. Pramod *et al*. (2014), in examining the sunflower EST database, found that microsatellites in coding regions were significantly shorter than those found in UTRs suggesting greater lability in UTR microsatellites.

We predicted that eSTRs are more likely to be found within genes that are constantly in need of “tuning” to respond to changes in the environment. Previous studies on bacterial species reported evidence in line with this prediction. Microsatellites were linked to the activity of some hypervariable regions regulating virulence at the interface of host pathogen interactions. These loci were fittingly named “contingency loci” (Moxon et al. 1994; Moxon *et al*, 2006). To test whether eSTRs are found in genes that are likely to show higher levels of evolutionary lability more so than others, we searched for specific GO terms enriched within eSTR-containing transcripts. The enrichment of GO terms linked to “regulation of transcription” (GO:0006355), “transcription factor activity” (GO:0003700), “DNA-binding” (GO:0003677), and “positive regulation of transferase activity” (GO:0051347) provide strong support for both cis-and trans-regulatory roles for eSTRs. Enriched GO terms such as, “spliceosomal complex” (GO:0005681), “mRNA splicing, via spliceosome” (GO:0000398), and “spliceosomal tri-snRNP complex” (GO:0097526) hint at specific mechanisms in which the involvement of eSTRs may be crucial (Table 1; Figure 5). Other more general GO terms identified as enriched within the eSTR-containing transcripts (Supplemental Table S15) also indicate specific genes where environment tracking and “tuning” may be desired.

Our study identified 479 transcribed microsatellites that can potentially serve as tuning knobs in common sunflower. Given that our study was limited to populations across a narrow latitudinal range, the number of microsatellites that show a significant effect on gene expression is noteworthy. Based on these findings, we envision that the number of microsatellites that could potentially alter phenotypes may be more than we could discover given the limited number of microsatellites investigated in this study. Our results based on transcribed regions suggest the existence of a substantial number of functional microsatellites that could potentially be under selection and may indicate the need to exercise caution when using transcribed microsatellites as neutral molecular markers in population genetic studies. Further, the results presented in this study consistent with the proposed functionality of microsatellites indicate the potential to predict a range of phenotypes based on specific microsatellite genotypes. This study provides strong evidence to suggest that microsatellites can rapidly generate heritable genetic variation which improves our understanding of mechanisms that can influence rapid adaptation via mutation at rates far greater than typically assumed.

## Author contributions

CR, GLW, MEC, and SP conducted the common garden experiment and collected data. MEC validated RNA-Seq based gene expression estimates with qRT-PCR. ADP oversaw the RNA Seq experiment. GLW, SP, and CR conducted the transcriptome analyses. MEW conceived and oversaw the study. CR performed the data analyses and wrote the manuscript. All authors read, revised, and approved the manuscript.

## Acknowledgements

This work was supported by the National Science Foundation grants, MCB- 1158521 to M.E.W and EPS-0903787 to A.D.P, and the Department of Biological Sciences at Mississippi State University. The authors wish to thank Lisa E. Wallace for specimen voucher preparation, Jessica L. Martin Judson for help with sample collection, and Brian S. Baldwin and Jesse I. Morrison for assistance with the common garden experiment.

## Data availability

All sequence data have been deposited at the National Center for Biotechnology Information short read archive under project PRJNA408292 and supplementary material has been deposited at figshare.

